# No evidence for association with APOL1 kidney disease risk alleles and Human African Trypanosomiasis in two Ugandan populations

**DOI:** 10.1101/180679

**Authors:** Magambo Phillip Kimuda, Harry Noyes, Julius Mulindwa, John Enyaru, Vincent Pius Alibu, Issa Sidibe, Dieuodonne Mumba, Christiane Hertz-Fowler, Annette MacLeod, Özlem Tastan Bishop, Enock Matovu (for the TrypanoGEN Research Group, as members of The H3Africa Consortium)

## Abstract

**Background:** Human African trypanosomiasis (HAT) manifests as an acute form caused by *Trypanosoma brucei rhodesiense* (Tbr) and a chronic form caused by *Trypanosoma brucei gambiense* (Tbg). Previous studies have suggested a host genetic role in infection outcomes, particularly for *APOL1*. We have undertaken a candidate gene association studies (CGAS) in a Ugandan Tbr and a Tbg HAT endemic area, to determine whether polymorphisms in *IL10, IL8, IL4, HLAG, TNFA, TNX4LB, IL6, IFNG, MIF, APOL1, HLAA, IL1B, IL4R, IL12B, IL12R, HP, HPR,* and *CFH* have a role in HAT.

**Methodology and results:** We included 238 and 202 participants from the Busoga Tbr and Northwest Uganda Tbg endemic areas respectively. Single Nucleotide Polymorphism (SNP) genotype data were analysed in the CGAS. The study was powered to find odds ratios > 2 but association testing of the SNPs with HAT yielded no positive associations i.e. none significant after correction for multiple testing. However there was strong evidence for no association with Tbr HAT and APOL1 G2 of the size previously reported in the Kabermaido district.

**Conclusions/significance:** A recent study in the Soroti and Kaberamaido focus in Central Uganda found that the APOL1 G2 allele was strongly associated with protection against Tbr HAT (odds ratio = 0.2). However, in our study no effect of G2 on Tbr HAT was found, despite being well powered to find a similar sized effect. It is possible that the G2 allele is protective from Tbr in the Soroti/Kabermaido focus but not in the Iganga district of Busoga, which differ in ethnicity and infection history. Mechanisms underlying HAT infection outcome and virulence are complex and might differ between populations, and likely involve several host, parasite or even environmental factors.

**Author Summary:** Human African Trypanosomiasis (HAT) occurs in two distinct disease forms; the acute form and the chronic form which are caused by microscopically indistinguishable hemo-parasites, *Trypanosoma brucei rhodesiense* and *Trypanosoma brucei gambiense* respectively. Uganda is the only country where both forms of the disease are found, though in geographically distinct areas. Recent studies have shown that host genetic factors play a role in HAT resistance and/or susceptibility, particularly by genes involved in the immune response. In this study, we identified single nucleotide polymorphisms in selected genes involved in immune responses and carried out a case-control candidate gene association study in Ugandan participants from the two endemic areas. We were unable to detect any polymorphisms that were robustly associated with either Tbr or Tbg HAT. However, our findings differ from recent studies carried out in the Tbr HAT another endemic area of Uganda that showed the APOL1 (Apolipoprotein 1) G2 allele to be protective against the disease which merits further investigation. Larger studies such as genome wide association studies (GWAS) by the TrypanoGEN network that has >3000 cases and controls covering seven countries (Cameroon, Cote d’Ivoire, DRC, Malawi, Uganda, Zambia) using the H3Africa customized chip reflective of African genetic diversity will present novel association targets.

## Introduction

The tsetse transmitted African trypanosomes are flagellated protozoa, a range of which cause disease in animals (known as Nagana) and humans (Human African Trypanosomiasis, HAT, also known as sleeping sickness). These diseases are responsible for significant morbidity and mortality [1–3] and therefore directly impact on public health and animal productivity. Current reports indicate that annual HAT incidence is on the decline, although under reporting is typical, especially in areas where conflicts and civil unrest interrupt control efforts and regular epidemiological surveys [4–6].

HAT is caused by two microscopically indistinguishable sub-species: *Trypanosoma brucei rhodesiense* that causes an acute form of the diseases that develops within a few weeks or months of infection, and *Trypanosoma brucei gambiense* that causes a chronic form of the disease that can take years to become patent. The acute form of the disease is prevalent in Eastern and Southern Africa while the chronic form of the disease is prevalent in West and Central Africa [4]. Uganda is the only country with active foci for both forms of the disease, though in geographically distinct regions.

Studies in the Democratic Republic of Congo (DRC), Cote D’Ivoire, Guinea and Uganda have found evidence for polymorphisms in IL6 and APOL1 associated with outcome of infection [7–9]. In the present study, we investigated the possible association of selected gene polymorphisms with HAT by undertaking a candidate gene association study (CGAS) using case-control samples from the Tbr and Tbg HAT endemic areas of Uganda. The *IL10, IL8, IL4, HLAG, TNFA, TNX4LB, IL6, IFNG, MIF, APOL1, HLAA, IL1B, IL4R, IL12B, IL12R, HP, HPR,* and *CFH* genes that were selected have protein products that are involved in the HAT immune response. The CGAS approach was used to compare the frequencies of genetic polymorphisms between cases and controls in order to identify risk variants for HAT in the two Ugandan populations.

## Materials and Methods

### Ethics and study population

This study was approved by the Uganda National Council of Science (UNCST; assigned code HS 1344) following review by the IRB of the Ministry of Health. Participants were identified through community engagement and active field surveys; they gave written informed consent administered in their local language by trained local health workers. In instances where participants were below 18 years of age, consent was sought from a parent or primary guardian. Any individuals for whom it was not possible to obtain consent or blood samples were excluded from the study.

The Tbr HAT endemic area samples were from the traditional Tbr HAT foci in the South East of Uganda [10]. Samples were collected mainly from Iganga district and included individuals from the predominantly Basoga ethnic group, with a few Baganda, Banyole, Balamogi, Basiginyi, Itesot, and Japadhola ethnicities.

The Tbg HAT endemic area samples were from the traditional Tbg HAT foci in the Northwest of Uganda [10]. Samples were collected from Adjumani, Arua, Koboko, Maracha, and Moyo districts and comprised of individuals from the Kakwa, Lubgbara and Madi ethnicities. In both areas, only individuals who were born and lived in these traditional foci were selected, as they were most likely exposed to HAT for most of their lives.

HAT cases were defined as individuals in whom trypanosomes have been detected in at least one of the body tissues including, blood, lymph node aspirates or cerebral spinal fluids. Controls were defined as individuals from the endemic area with no history or any signs/symptoms suggestive of HAT. Controls from the Tbg HAT endemic area were required to have no serological reaction to the CATT or Trypanolysis test.

Blood was drawn by venipuncture and collected in EDTA/heparin vacutainer tubes (BD). Buffy coats were prepared from the whole blood in field laboratories using centrifugation, aliquoted, and then stored in liquid nitrogen in preparation for DNA extraction that was carried out at the Molecular Biology Laboratory, COVAB, Makerere University. The DNA was quantified using a Qubit™ (Life Technologies).

### Study design

This study was one of five studies of populations of HAT endemic areas in Cameroon, Cote d’Ivoire, Guinea, Malawi and Uganda. The studies were designed to have 80% power to detect odds ratios (OR) >2 for loci with disease allele frequencies of 0.15 – 0.65 and 100 cases and 100 controls with the 96 SNPs genotyped. The study design included an overall total of 462 samples, 239 samples from Tbr HAT endemic regions (120 cases, 119 controls) and 223 samples from Tbg HAT endemic regions (110 cases and 113 controls).

Power calculations were undertaken using the pbsize routine in Genetics Analysis Package gap version 1.1-16 in R [11].

### Gene Selection

The selection of the genes depended on prior knowledge of the genes and their association with the HAT. The following genes *IL10* [9]*, IL8* [7]*, IL4* [12]*, HLAG* [13]*, TNFA* [7]*, TNX4LB* [14]*, IL6* [7]*, IFNG* [15]*, MIF* [16]*, APOL1* [8], *HLAA* [17]*, IL1B* [18]*, IL4R* [18]*, IL12B* [18]*, IL12R* [18]*, HP* [19]*, HPR* [19,20], and *CFH* [21] were selected.

### SNP Selection

96 SNP were selected for genotyping using two strategies: 1) SNP that had been previously reported to be associated with HAT or 2) in the cases of *IL4, IL8, IL6, HLAG* and *IFNG* by complete scans with linked marker SNP (r^2^ < 0.5) across each gene. The SNPs in this second group of genes were selected using a merged SNP dataset obtained from 10X coverage whole genome sequence data generated from 230 residents living in regions (DRC, Guinea Conakry, Ivory Coast and Uganda) where trypanosomiasis is endemic (TrypanoGEN consortium, sequences at European Nucleotide Archive Study: EGAS00001002482) and 1000 Genomes Project data from African populations. Linkage (r^2^) between loci was estimated using Plink [22] and sets of SNPs that covered the gene were identified. Some SNP loci were excluded during assay development or failed to genotype and were not replaced.

### Genotyping

Approximately 1μg of gDNA per sample were submitted to INRA (Plateforme Genome Transcriptome de Bordeaux, France) for genotyping. A multiplex analysis (two sets of 80 SNPs each) was designed using Assay Design Suite v2.0 (Agena Biosciences). SNP genotyping was achieved with the iPLEX Gold genotyping kit (Agena Biosciences) for the MassArray iPLEX genotyping assay, following the manufacturer’s instructions. Products were detected on a MassArray mass spectrophotometer and the data acquired in real time with MassArray RT software (Agena Biosciences). SNP clustering and validation was carried out with Typer 4.0 software (Agena Biosciences). SNPs that failed genotyping at INRA and some additional SNPs were genotyped at LGC Genomics, Hoddesden, UK where SNP were genotyped using the PCR based KASP assay [23]. A summary of the candidate genes and SNPs is shown in Supplementary Table 1.

### Statistical analysis

The raw genotypic data were converted to PLINK format and quality control (QC) procedures implemented using the PLINK v1.9 package (http://pngu.mgh.harvard.edu/purcell/plink/) [22]. PLINK was used to determine the level of individual and genotype missingness, Hardy-Weinberg Equilibrium (HWE), estimate allele frequencies, and linkage disequilibrium (LD). Testing for population stratification and admixture was carried out using Admixture 1.3 [24] and the plot was visualized using StructurePlot2 [25].

Testing for the association of SNPs with HAT was done using a Fisher’s exact test [40] implemented in PLINK at 95% confidence level. Controlling for multiple testing was implemented using a Bonferroni correction (α* = α/n, where α* is the corrected *P*-value, α is the level of significance and n is the number of independent SNP association tests) [27]. The Bonferroni correction assumes that each of the statistical tests are independent; however, this was not always true since there was some linkage disequilibrium between the SNPs in *IL4, IL8, IL6, HLAG* and *IFNG* which were subject to complete linkage scans. Where the assumption of independence is not true, the correction is too strict potentially leading to false negatives. Thus, a less stringent correction for multiple testing was also employed. The Benjamini-Hochberg false discovery rate (FDR) estimates the proportion of significant results (*P* < 0.05) that are false positives [27,28].

## Results

Our study population consisted of 239 individuals from Tbr and 223 from the Tbg HAT endemic areas. The former comprised of 120 cases and 119 controls, who had a mean age of 43 ± 5 years, and a male to female ratio of 1:2. The Tbg HAT endemic area participants comprised of 110 cases and 113 controls, who had a mean age of 37 ± 5 years, and a male to female ratio of 1:1.

### Genotyping and data quality control

Ninety-six (96) SNPs in 15 genes were genotyped from each of the Tbr and Tbg HAT endemic area samples as shown in supplementary table 1 (the Plink MAP and PED files are available in Supplementary data 1-3). Before association testing, individuals with missing data, SNPs that were not in HWE, SNPs with missing data or those that were poorly genotyped were removed using PLINK [22,29].

Individuals with more than 20% or 15% missing data were excluded from the Tbr and the Tbg HAT endemic datasets, respectively, resulting in a final dataset of 238 (119 cases and 119 controls, 1:2 male to female sex ratio) individuals from the Tbr HAT endemic sample and 202 (99 cases and 103 controls, 1:1 male to female sex ratio) individuals from the Tbg HAT endemic sample (Supplementary Figures 1-2). Similarly, SNPs that were missing more than 30% or 40% data were excluded from the Tbr and the Tbg HAT endemic area samples (Supplementary Figures 3-4). We used a HWE p-value cut-off of 1 x 10^-8^ and further selection of SNPs below the HWE cut off was done basing on their genotype scatter plots to see which loci were to be excluded. Furthermore, SNPs that were in a five SNP window after a single step with a variance inflation factor (VIF) [VIF = 1/(1-R^2^)] beyond 0.2 were excluded from both sample datasets. After quality pruning, 79 SNPs from Tbr and 85 SNPs from the Tbg HAT endemic samples were included in the association testing.

### Admixture for population structure

Admixture was used to test for population structure that might confound the association study. Eight values of *K* ancestral populations from 1-8 were tested to identify which had the lowest coefficient of variations (CV) error. CV error was at a minimum for *K*=4, but the CV error was very similar for all values of *K* (0.42 - 0.46) providing no persuasive evidence for any particular number of ancestral populations. The Admixture plot showed no clear evidence for any gross population structure and therefore no correction for population structure was applied in the analysis.

### Association testing yielded no robust associations

Five SNPs in the Tbr HAT endemic area and four in the Tbg endemic had raw p < 0.05 but none of these remained significant after Bonferroni correction (Table 1). Surprisingly, there was no evidence for association with any SNP in APOL1.

**Table 1:** SNPs that showed the lowest *p* values after association testing with Tbr and Tbg HAT.

| Tbr HAT endemic sample (N= 238; 119 cases, 119 controls) |  |  |  |  |  |  |  |  |  |  |  |  |  |
| --- | --- | --- | --- | --- | --- | --- | --- | --- | --- | --- | --- | --- | --- |
| CHR | SNP | GENE | BP | Allele 1 | Cases | Controls | Allele 2 | P | OR | 95%<br>upper CI | 95 %<br>lower CI | BONF | FDR_BH |
| 6 | rs9380142 | HLA-G | 29798794 | G | 0.369 | 0.242 | A | 0.003 | 1.834 | 1.231 | 2.731 | 0.1805 | 0.181 |
| 6 | rs1233330 | HLA-G | 297991036 | A | 0.076 | 0.136 | G | 0.03 | 0.522 | 0.3419 | 1.019 | 1 | 0.434 |
| 5 | rs2243283 | IL4 | 132016593 | G | 0.275 | 0.188 | C | 0.04 | 1.644 | 1.031 | 2.621 | 1 | 0.434 |
| 22 | rs34383331 | MIF | 24238079 | A | 0.24 | 0.16 | T | 0.03 | 1.657 | 1.031 | 2.621 | 1 | 0.434 |
| 22 | rs9282783 | MIF | 24236359 | G | 0.089 | 0.042 | C | 0.033 | 2.227 | 1.031 | 2.621 | 1 | 0.434 |
| 6 | rs1800630 | TNFA | 31542476 | A | 0.156 | 0.092 | C | 0.038 | 1.807 | 1.031 | 3.169 | 1 | 0.434 |
| Tbg HAT endemic sample (N=202; 99 cases, 103 controls) |  |  |  |  |  |  |  |  |  |  |  |  |  |
| CHR | SNP | GENE | BP | Allele 1 | Cases | Controls | Allele 2 | P | OR | 95%<br>upper CI | 95 %<br>lower CI | BONF | FDR_BH |
| 1 | rs1061170 | CFH | 196659237 | C | 0.409 | 0.525 | T | 0.019 | 0.627 | 0.4221 | 0.9313 | 1 | 0.611 |
| 6 | rs1233330 | HLA-G | 29799103 | A | 0.076 | 0.136 | G | 0.045 | 0.521 | 0.5211 | 0.2693 | 1 | 0.611 |
| 12 | rs78554979 | IFNG | 68554636 | C | 0.051 | 0.015 | T | 0.035 | 3.638 | 0.9861 | 13.42 | 1 | 0.611 |
| 7 | rs2069843 | IL6 | 22769994 | A | 0.147 | 0.078 | G | 0.033 | 2.038 | 1.07 | 3.882 | 1 | 0.611 |

\*Abbreviations: CHR = Chromosome, SNP = SNP ID, BP = Physical position (base-pair) (Human genome build GRCh37), Allele 1 = Minor allele name (based on whole sample), Cases = Frequency of this allele in cases, Controls = Frequency of this allele in controls, Allele 2 = Major allele name, P = Asymptotic p-value for this test, OR = Estimated odds ratio (for Allele 1, i.e. Allele 2 is reference), BONF = Bonferroni single-step adjusted p-values, FDR\_BH = Benjamini & Hochberg (1995) step-up FDR control.

## Discussion

In this case-control CGAS, we found no robust evidence for variants associated with Tbr and Tbg HAT in two Ugandan populations. We tested for association between candidate genes and the disease caused by Tbg and Tbr separately as they present two distinct forms of the disease. Tbr and Tbg parasite resistance to human serum is mediated by different mechanisms which place distinct selective pressures on the host genes [30]. Furthermore, the two populations were from different broad ethnolinguistic groups, and were geographically isolated from each other [10]. Admixture analysis found no evidence of population structure with these SNP which might have reduced the power of the study (Supplementary Figure 5). We found no SNP associated with HAT after multiple testing corrections. Our power calculations indicated that we had power to detect odds ratios > 2, however 7 of the 10 SNP with P <0.05 had odds ratios < 2.0, which the study was not powered to detect. Larger populations would be required to confirm these populations and the data presented could be used to estimate the necessary sample size.

The most striking feature of the data was the absence of any association at *APOL1*. The APOL1 G2 (rs71785313) allele has been shown to be lytic to *T. b. rhodesiense in vitro* [31] and a recent study in the Soroti and Kaberamaido focus in Eastern Uganda found an association with APOL1 G2 and protection from Tbr HAT with an odds ratio of 0.2 [8]. The present study in the Busoga focus was well powered to discover such a strong effect, but the frequencies of *APOL1* G2 in cases and controls was almost equal (8.1% and 8.6%) with a 95% confidence interval for the odds ratio of (0.37-2.34) indication that an odds ratio as large as seen in Kabermaido is very unlikely to be seen in Busoga (Supplementary data Table S2). Another TrypanoGEN study in a *T. b. rhodesiense* endemic area of Malawi has also found no association with G2 despite higher frequencies of the protective allele (14%) (Kelita 2017) [Submitted to PLOS NTD]. Therefore, despite the well-established function of APOL1 in response to trypanosome infection and the evidence for protection associated with G2 in Kabermaido [8], the role of APOL1 G2 in response to *T. b. rhodesiense* infection more generally remains to be clarified.

In conclusion, despite the suggestively significant associations found at nine SNP loci, none of them passed Bonferroni correction for multiple testing [27]. FDR_BH indicated that there was a greater than 5% probability for each of these SNPs being associated with HAT [27,28]. The finding of suggestive associations in multiple populations would increase the probability that these are genuine associations with disease [32]. For example, our findings suggest that *HLA-G* variants may be important in both forms of the disease. These observations will be followed up by the TrypanoGEN network which has collected >3,000 cases and controls from seven regions in six countries (Cameroon, Cote d’Ivoire, DRC, Malawi, Uganda, Zambia) [33]. The samples will be genotyped using the H3Africa customized SNP chip that is reflective of the diversity within ethnolinguistic groups in Africa, presently under development the H3A consortium.

## Acknowledgments

We are grateful for the support and participation of the study populations included in this study, as well as the technical team at COVAB and the participating hospitals who executed the field surveys.

## SUPPORTING INFORMATION

**S1 Fig:**
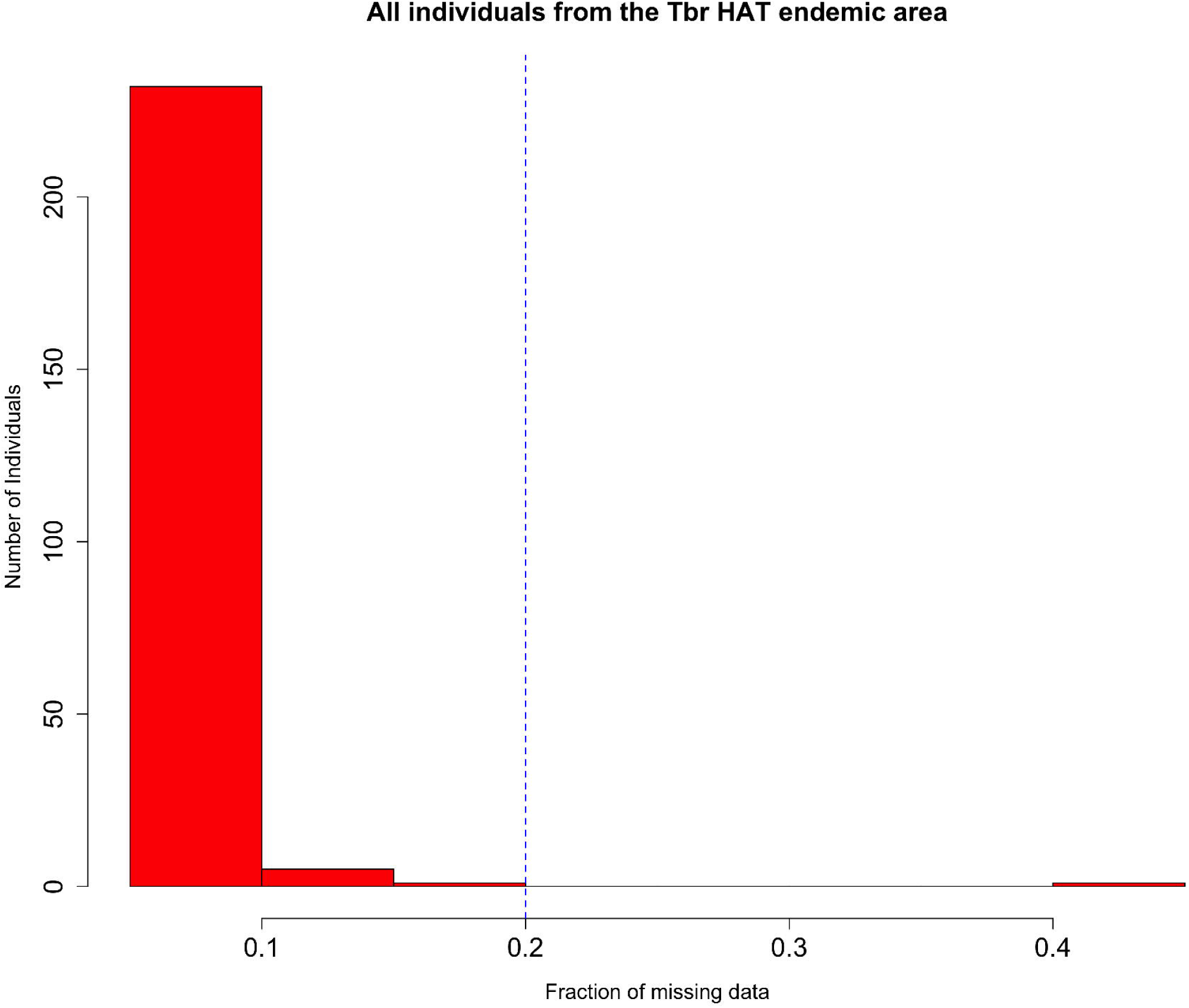
Histogram of missing data rate in all individuals from the Tbr HAT endemic area. The dashed vertical line represents a 20% threshold used in the exclusion criteria due to excessive failure rate.

**S2 Fig:**
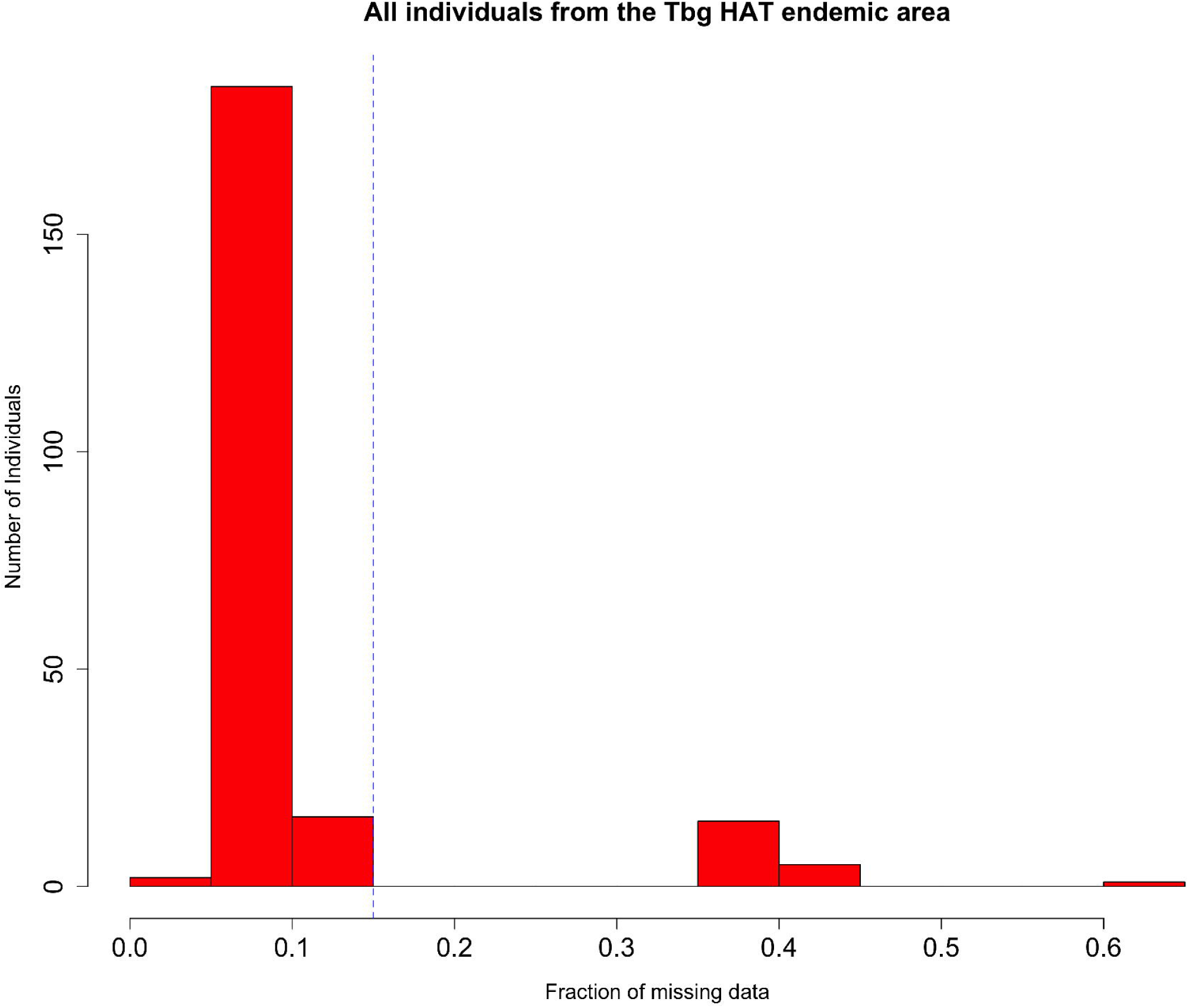
Histogram of missing data rate in all individuals from the Tbg HAT endemic area. The dashed vertical line represents a 15% threshold used in the exclusion criteria due to excessive failure rate.

**S3 Fig:**
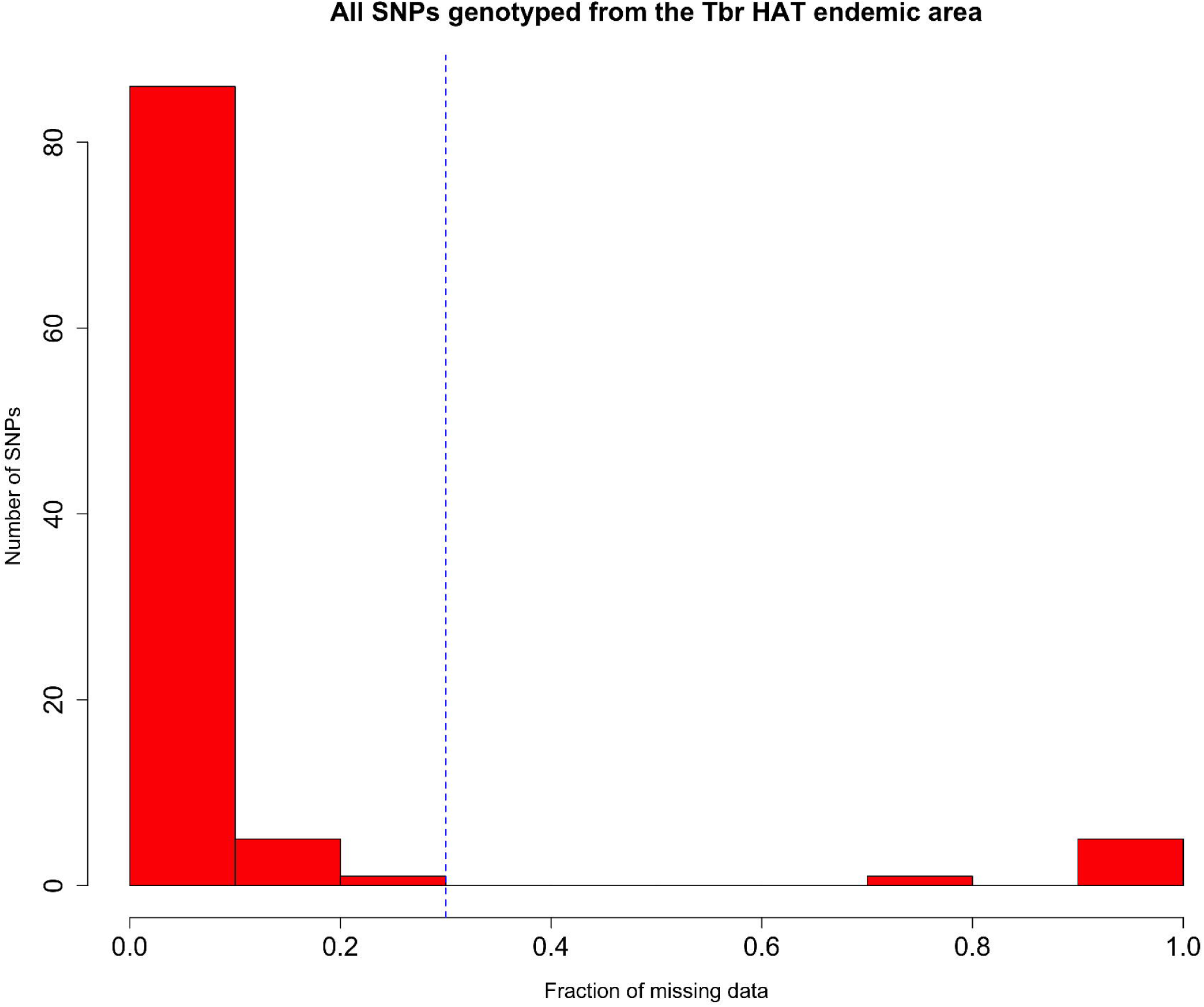
Histogram of missing data rate in all SNPs from the Tbr HAT endemic area passing. The dashed vertical line represents a 30% threshold used in the exclusion criteria due excessive failure rate.

**S4 Fig:**
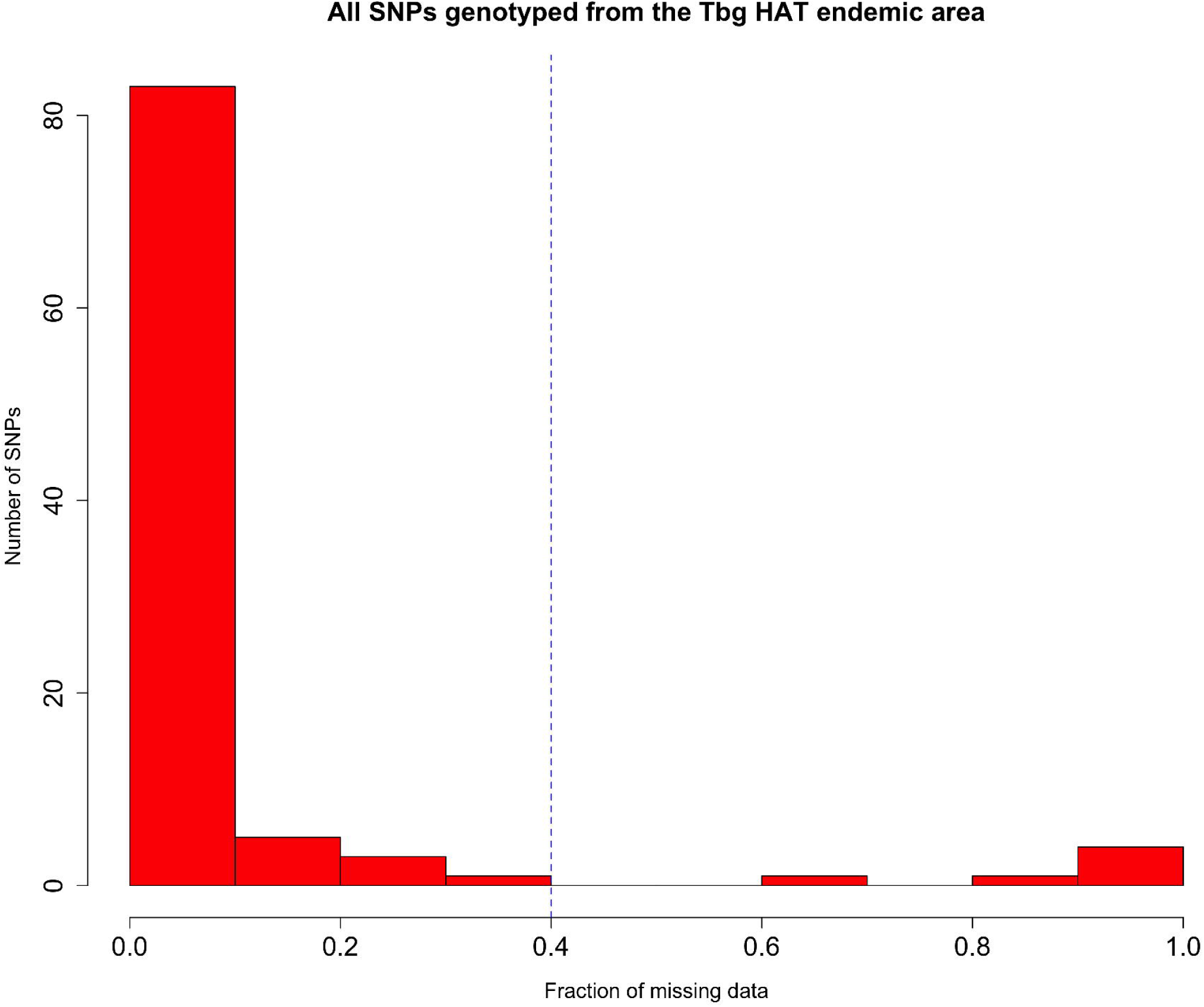
Histogram of missing data rate in all SNPs from the Tbg HAT endemic area passing. The dashed vertical line represents a 40% threshold used in the exclusion criteria due to excessive failure rate.

**S5 Fig:**
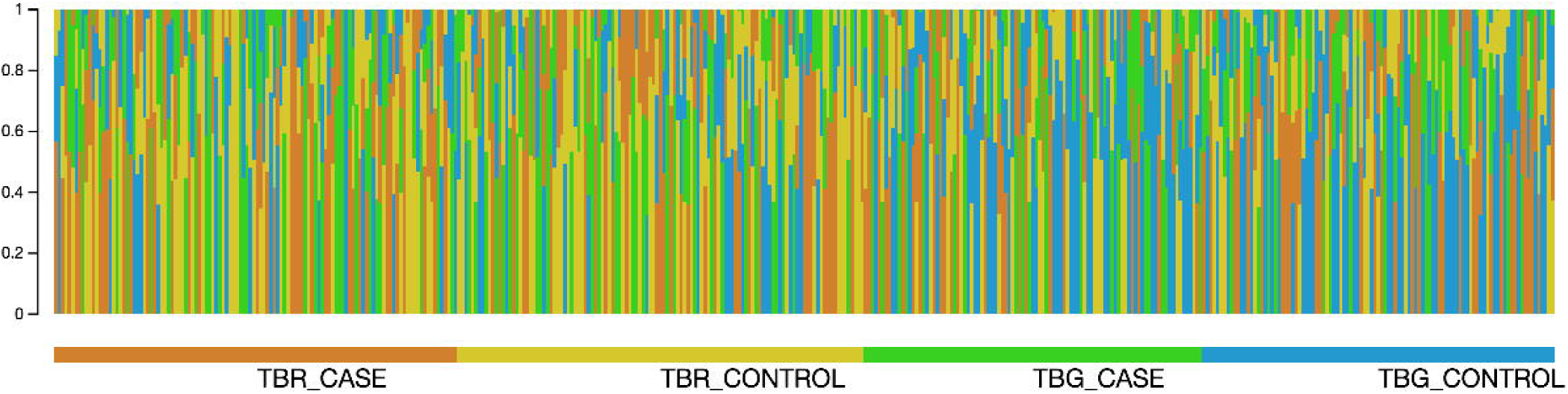
Bar plot showing the admixture analysis performed for K=4. Individuals are shown as vertical bars.

**S1 Table: Candidate genes included in the study.**

**S2 Table: Association results of 65 SNPs with Acute HAT.** *Abbreviations: CHR = Chromosome, SNP = SNP ID, BP = Physical position (base-pair), A1 = Minor allele name (based on whole sample), F_A = Frequency of this allele in cases, F_U = Frequency of this allele in controls, A2 = Major allele name, P = Asymptotic p-value for this test, OR = Estimated odds ratio (for A1, i.e. A2 is reference), BONF = Bonferroni single-step adjusted p-values, FDR_BH = Benjamini & Hochberg (1995) step-up FDR control, FST = Fixation index, and MAF = Minor allele frequency. The level of significance is 0.05.

**S3 Table: Association results of 65 SNPs with Chronic HAT.** *Abbreviations: CHR = Chromosome, SNP = SNP ID, BP = Physical position (base-pair), A1 = Minor allele name (based on whole sample), F_A = Frequency of this allele in cases, F_U = Frequency of this allele in controls, A2 = Major allele name, P = Asymptotic p-value for this test, OR = Estimated odds ratio (for A1, i.e. A2 is reference), BONF = Bonferroni single-step adjusted p-values, FDR_BH = Benjamini & Hochberg (1995) step-up FDR control, FST = Fixation index, and MAF = Minor allele frequency. The level of significance is 0.05.

**S1 DATA**: A read me text with a brief description of the TrypanoGEN data.

**S2 DATA**: The TrypanoGEN Uganda samples MAP file.

**S3 DATA**: The TrypanoGEN Uganda samples PED file.

